# Acidic Sphingomyelinase Interactions with Lysosomal Membranes and Cation Amphiphilic Drugs: a Molecular Dynamics Investigation

**DOI:** 10.1101/2023.12.14.571676

**Authors:** Simone Scrima, Matteo Lambrughi, Kenji Maeda, Marja Jäättelä, Elena Papaleo

**Affiliations:** Cancer Structural Biology, Danish Cancer Institute, 2100, Copenhagen, Denmark; Cancer System Biology, Section for Bioinformatics, Department of Health and Technology, Technical University of Denmark 2800, Lyngby, Denmark; Cell Death and Metabolism, Center for Autophagy, Recycling and Disease, Danish Cancer Institute, 2100, Copenhagen, Denmark; Department of Cellular and Molecular Medicine, Faculty of Health Sciences, University of Copenhagen, 2200, Copenhagen, Denmark

## Abstract

Lysosomes are pivotal in cellular functions and disease, influencing cancer progression and therapy resistance with Acid Sphingomyelinase (ASM) governing their membrane integrity. Moreover, cation amphiphilic drugs (CADs) are known as ASM inhibitors and have anti-cancer activity, but the structural mechanisms of their interactions with the lysosomal membrane and ASM are poorly explored.

Our study, leveraging all-atom explicit solvent molecular dynamics simulations, delves into the interaction of glycosylated ASM with the lysosomal membrane and the effects of one of the CAD representatives, i.e., ebastine on the membrane and ASM.

Our results confirm the ASM association to the membrane through the saposin domain, previously only showed with coarse grained models. Furthermore, we elucidated the role of specific residues and ASM-induced membrane curvature in lipid recruitment and orientation. Ebastine also interferes with the association of ASM with the membrane at the level of a loop in the catalytic domain engaging in membrane interactions. Our computational approach, applicable to various CADs or membrane compositions, provides insights into ASM and CAD interaction with the membrane, offering a valuable tool for future studies.

## Introduction

Lysosomes are the cellular organelles known as the cell’s recycling centers and key elements in preserving cellular energy homeostasis ^1^. Furthermore, the cellular roles of lysosomes extend beyond degradation and recycling, as they are involved in a broad range of processes, including cell death, metabolic adaptation, and antigen presentation ^2,3^, and in disease, as in cancer where they are involved in promoting cell growth, invasion, metastasis and drug resistance ^4^. In particular, lysosomal membrane permeabilization and leakage (i.e., the release of lysosomal contents into the cytosol) can trigger cell death pathways (i.e., lysosome-dependent cell death) ^5,6^. Nevertheless, it has been shown that the lysosomal membrane integrity is dynamically regulated in a widespread range of critical physiological cellular processes ^3,7,8^ The stability of the lysosomal membrane is ensured by a protective glycocalyx and controlled by numerous lysosomal enzymes, including acid sphingomyelinase (ASM). ASM is a lysosomal and peripheral membrane phosphodiesterase that catalyzes the hydrolysis of sphingomyelins to ceramide and phosphocholine ^9,10^.

The structure of ASM includes different domains, i.e., a saposin-like domain (residues 86-169), a proline-rich linker (residues 170-197) that connects the saposin domain to the catalytic domain (residues 198-540) (**Figure 1A**). Moreover, there is a helical domain at the C-terminus (residues 541-613). The mature form of the protein includes six N-linked glycosylation sites ^11–13^, and the protein is primarily active as a monomer ^12^. ASM coordinates two zinc ions with a trigonal bipyramidal coordination geometry thanks to two aspartic acids (D208, D280), four histidines (H210, H427, H459, H461), one asparagine (N320), and a catalytic water molecule ^11^ (**Figure 1A**). Moreover, three residues (i.e., N327, E390, and Y490) have been suggested for substrate binding.^11^ The active site geometry could favor proton donation to the oxyanion of the ceramide leaving group by H321 or by H284 ^11^ The active site of ASM presents a concave and “bowl-like shape” surface, including the saposin domain’s inner surface. It is oriented towards the membrane, where it recruits and binds sphingomyelin and other lipid substrates. It has been shown that ASM has a broad lipid substrate specificity in vitro and can process several membrane phospholipids, including ceramide-1-phosphate and bis(monoacylglycero)phosphate (BMPs).^9^ Thus, it has been proposed that ASM could play a key role in phospholipid catabolism, acting as a promiscuous phospholipase^9^. ASM is suggested to pH-dependently anchor at the membranes of the intralysosomal luminal vesicles, which are known to be the platform for the catabolism of lipids ^12^, being tethered by the presence of anionic phospholipids, such as BMPs ^14^. It has been suggested that the membrane anchor allows ASM activity. When the interaction with the membrane is lost or altered, the protein is rapidly inactivated and degraded by cathepsins ^6,14^. Nevertheless, the conformations and dynamics of ASM when associated with the lysosomal membrane, such as in the recruitment of lipids at the catalytic site, are unclear. The work of Xiong et al. ^12^ provided insight into the association of ASM to a lipid bilayer composed of anionic 1-palmitoyl-2-oleoyl-sn-glycero-3-phosphoglycerol lipids using μs coarse-grain molecular dynamics (MD) simulations. Xiong et al. performed coarse-grained MD simulations starting with ASM randomly oriented with respect to the lipid bilayer. The authors observed two lipid-binding modes involving the saposin domain: a) type I and b) type II ^12^. The type I association was characterized by the partial insertion of the saposin domain into the lipid headgroup region of the bilayer, where helices H1, H2, and H4 predominantly make contact with the lipids, and helix H3 makes only a few contacts. In contrast, the type II association presented a more open conformation of the saposin domain, allowing all four helices, H1-4, to form contacts with lipids ^12^.

**Figure 1.**
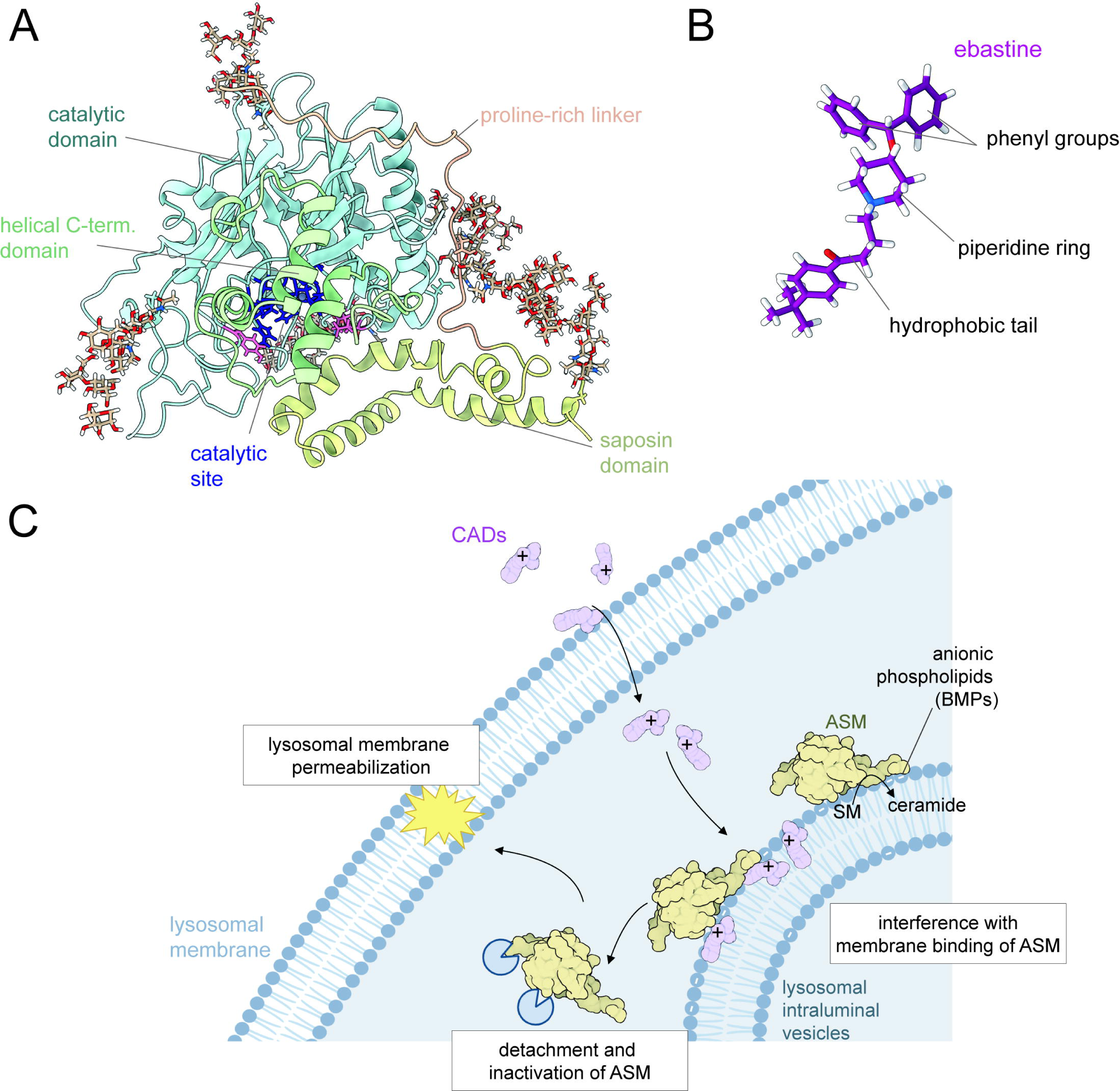
Mechanisms of ASM and cationic amphiphilic drugs. (A) Structure of human ASM as cartoon representation, highlighting its distinct domains: the saposin domain (residues 86-169, yellow), the proline-rich linker (residues 170-197, light brown), the catalytic domain (residues 198-540, light blue), and the helical domain at the C-terminus (residues 541-613, light green). The catalytic site of ASM, characterized by a concave shape, coordinates two zinc ions (depicted as black spheres). The zinc-coordination residues (D208, D280, H210, H427, H459, H461, N320) and the two residues that may facilitate proton donation in the catalytic mechanism are shown as blue sticks. Additionally, three residues (N327, E390, and Y490), suggested to be important for substrate binding, are represented as pink sticks. The N-glycosylation sites are illustrated as light brown sticks. (B) Structure of ebastine, a cationic amphiphilic drug (CAD). CADs typically possess a hydrophobic tail and one or more hydrophilic, basic groups, such as the tertiary amine group in the piperidine ring of ebastine. (C) Proposed anticancer mechanisms of CADs. The mechanism suggests that unprotonated CADs can passively diffuse across lysosomal membranes, becoming protonated and trapped inside the lysosomes. Here the positively charged CADs can insert in the lysosomal intraluminal vesicles and interact with anionic lipids like BMPs, crucial for ASM tethering. This could disrupt the binding of ASM to the membrane, leading to its dissociation and cathepsin-mediated degradation. The resulting accumulation of sphingomyelin and lysoglycerophospholipids may cause lysosomal membrane permeabilization, releasing cathepsins and cytotoxic contents into the cytosol, thus initiating lysosome-dependent cell death. The illustration has been created with BioRender.com.

Notably, ASM and lysosomes are targets of interest that can be potentially exploited in cancer therapy ^6,15,16^. Growing data reveal that cancer cells, especially in metastatic cancers, undergo alterations in their lysosomes, including changes in lipid composition, that, while allowing escaping from apoptosis induced by therapy, increase sensitivity to lysosome-dependent cell death ^17,18^. Clinically relevant drugs have been reported to induce lysosomal permeabilization and lysosome-dependent cell death in cancer cells ^19,20^. Several of them are cationic amphiphilic drugs (CADs), which have been reported to have multiple putative anti-cancer activities, including acting as functional inhibitors of ASM ^6,19,21–23^. Several studies reported the potential of CADs against cancer, showing their cancer-specific cytotoxicity against a wide range of cancer cells. Their effectiveness in animal cancer models and pharmaco-epidemiological studies suggested better outcomes for cancer patients taking CADs ^6,24,25^ CADs, such as ebastine, are a diverse group of pharmacological compounds used to treat various human conditions, including allergies, mental health disorders, cardiovascular diseases, and infections ^6^. CADs generally include a hydrophobic part, often made up of aromatic or aliphatic rings, and one or more hydrophilic and basic groups, such as amine groups (**Figure 1B**). The proposed mechanism of anticancer action of CADs suggests that due to their basic and amphiphilic nature, they can diffuse across the lysosomal membrane and then be protonated and trapped inside the lysosomes ^26–28^ (**Figure 1C**). The accumulation of CADs in the lumen of lysosomes increases the lysosomal pH ^22^ and CADs could insert into the membranes of the intraluminal lysosomal vesicles ^28^, which are the sites of lysosomal lipid degradation and to which ASM and other lysosomal lipases anchor to interact with their lipid substrates ^14^ (**Figure 1C**). Here, the presence of CADs could interact with anionic lipids required for the tethering of ASM to the lysosomal intraluminal vesicles, altering the binding of ASM to the membrane and allowing cathepsin-mediated degradation ^19,29^ (**Figure 1C**). The consequent accumulation of sphingomyelin and lysoglycerophospholipids induces lysosomal membrane permeabilization, resulting in the release of cathepsins and cytotoxic contents into the cytosol and initiating lysosome-dependent cell death ^21,30^. However, several steps of this mechanism are still unclear, including how and if CADs insert within the membranes and impact membrane properties and stability and how they affect ASM anchoring to the membrane and its lipid-protein interactions.

In this study, we used MD simulations to characterize the interaction between the fully glycosylated ASM form and the lysosomal membrane and to understand the effects of ebastine, as an example of CADs, on the lysosomal membrane and the ASM protein.

## Results and Discussion

### Analysis of lipidomics data of lysosomes of HeLa and U2OS cells

To study the binding of ASM to the membrane, we designed bilayers as models of lysosomal membranes by analyzing lipidomic datasets of lysosomes of HeLa and U2OS cells ^31,32^. The lysosomes were previously isolated by immunoaffinity purification using an antibody against the lysosomal-associated membrane protein 1 and profiled by performing quantitative mass spectrometry-based shotgun lipidomics analysis ^31–34^. More than 300 lipid species of 28 different lipid classes of the categories of glycerolipids, fatty acyls, glycerophospholipids, sphingolipids, and sterol lipids were quantified as molar percentages (mol%, molar quantity normalized relative to the total molar quantities of all the identified lipids) ^31,32^(**Figure 2A**). We compared these lipidomics data with the lipid compositions of a recent computational model of the membrane of mammalian lysosomes (**Figure 2A**) ^35^. The two cell lines showed very similar lipid compositions for their lysosomes (**Figure 2A**). Cholesterol, phosphatidylcholine and phosphatidylethanolamines are the most abundant lipids in all the three datasets. The main difference is associated with the higher percentage of cholesterol in both U2OS and HeLa cells, while phosphatidylethanolamines are overrepresented in the computational model (**Figure 2A**). On the other hand, the content of lysosome-specific BMP (phosphatidylglycerol-BMP) and sphingomyelin classes are consistent across the three datasets, accounting for around 2.3–2.8 mol% and 8-6 mol% of lipids in the lipidomics datasets and around 7% and 6% in the computational model, respectively (**Figure 2A**). When considering the lipidomic data, it is essential to note that the samples encompass both the external lysosomal membranes and the intralysosomal luminal vesicles (see Materials and Methods). BMPs localize primarily at the intralysosomal luminal vesicles where ASM anchors and plays its activity ^36^. Consequently, one would expect that the BMP mol% should be higher in lipidomics of purified intralysosomal luminal vesicles. Given the overall similar patterns of lipid compositions between the lipidomic dataset of lysosomes of HeLa and U2OS cells and the computational model, we designed our lipid bilayer to resemble the latter.

**Figure 2.**
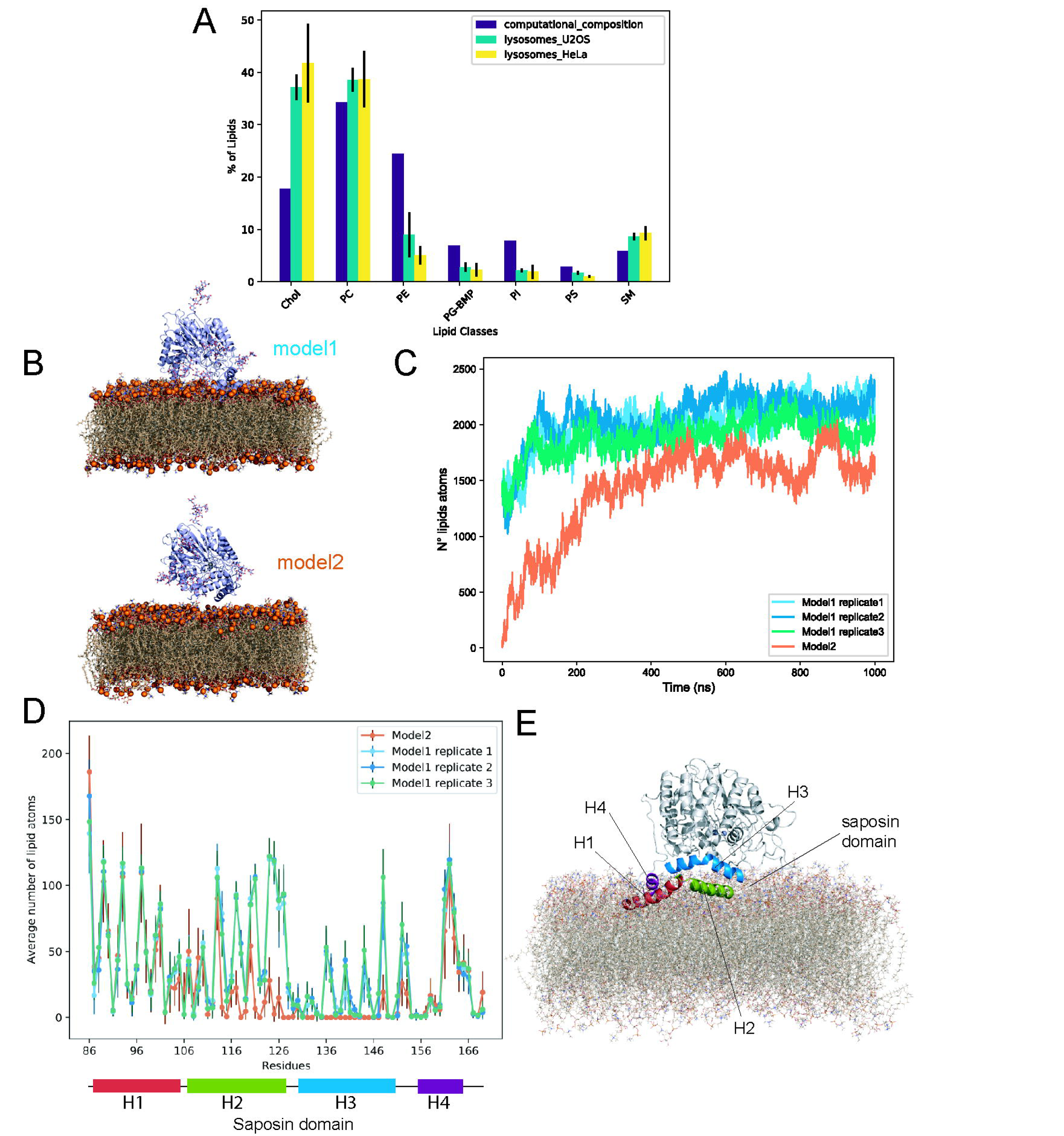
ASM interacts with lipid bilayers resembling lipid compositions of lysosomal membranes in all-atom MD simulations. (A) Mass spectrometry-based shotgun lipidomics profiles of immunopurified lysosomes from HeLa (yellow bars) and U2OS (green bars) cells, compared with a computational model of mammalian lysosomal membranes (purple bars). The two cell lines show very similar lipid compositions. The analysis reveals similar lipid compositions across the two cell lines and the model, consistent with lysosome-specific BMPs-phosphatidylglycerol (PG) and sphingomyelin (SM) classes. (B) Two modeling strategies were used to start the MD simulations: i) model 1, where ASM is positioned directly in contact with the bilayer surface (upper panel), and model 2, where ASM is initially distanced from the bilayer (lower panel). ASM is represented as a blue cartoon, and the lipids are shown as light brown sticks with phosphate groups in their headgroups highlighted as orange spheres. (C) The number of lipid atoms in contact with ASM during the MD simulations shows the spontaneous association of ASM in both model 1 and model 2. (D) The average number of lipid atoms in contact with the saposin domain of ASM indicates a partial insertion of this domain into the bilayer. Helices H1, H2, and H4 are predominantly involved in the interaction with the bilayer. (E) An example of the orientation of ASM when in contact with its saposin domain to the lipids was observed from replicate 1 of model 1.

### Interaction between ASM and the lysosomal membrane

To characterize the interaction between ASM and the lysosomal membrane, we collected four one-μs all-atom MD simulations of the protein with the lipid bilayer **(Table1)**, employing two strategies to predict how they associate **(Figure2B)**. A model and its first MD simulation (i.e., replicate 1 of model 1) are taken from ref ^37^. This initial system for MD simulations had ASM positioned on the bilayer surface and interacting with the lipids. We performed two additional MD replicates for this configuration (**Table 1**). Secondly, we carried out one additional MD simulation, positioning the protein 15 Å away from the bilayer, along the axis perpendicular to the surface of the bilayer. This was done to avoid extensive contact, following a previously suggested approach ^38^. We will refer hereinafter to model 1 and model 2 (**Figure 2B**) for the two different initial configurations, respectively. In addition, we accounted for the N-glycosylations of ASM in the interaction with the membrane. Hence, we modeled a fully Man5-glycosylated variant of ASM (i.e., at the sites N88, N177, N337, N397, N505, and N522), as previously described ^37^.

**Table 1.** Summary of the molecular dynamics simulations included in the study.

| System | Starting distance of protein from the bilayer (Z-axis, Å) | Temperature (K) | Headgroup molar ratio | N° lipids | Simulation time ( μs) | N° replicates |
| --- | --- | --- | --- | --- | --- | --- |
| Model 1 (ASM<sub>1 86-613</sub>) | 0 | 310 | PC:PE:PI:PS:SM:CH<br>OL:BMP(35:25:8:3:6:18:7) | 612 | 1 | 3 |
| Model 2 (ASM<sub>2 86-613</sub>) | 15 | 310 | PC:PE:PI:PS:SM:CH<br>OL:BMP(35:25:8:3:6:18:7) | 612 | 1 | 1 |
| EBAH-ASM Model 1 + ebastine (ASM<sub>1 86-613</sub>) | 0 | 310 | PC:PE:PI:PS:SM:CH<br>OL:BMP(35:25:8:3:6:18:7) | 612 | 0.5 | 3 |
| EBAH bilayer + ebastine | N.A. | 310 | PC:PE:PI:PS:SM:CH<br>OL:BMP(35:25:8:3:6:18:7) | 612 | 0.5 | 3 |

We investigated the structural properties of the lipid bilayer and the effects induced by the association with ASM. We noticed that the four simulations have a comparable average lipid density of the bilayer (∼ 13000 Å^-3^) (**Figure S1** and OSF repository). However, we observed that in the MD simulation starting from model 1 (i.e., already localized on top of the membrane at the beginning of the simulations), there is a higher density of lipids in several areas of the bilayer, suggesting that they are more ordered and packed (**Figure S1**). Furthermore, we calculated the average area-per-lipid and lipid bilayer thickness in the four MD simulations. We observed that MD simulations of model 1 and model 2 have similar values for both area-per-lipid (average 54.0 Å^2^ and 53.5 Å^2^, respectively) and thickness (average 40.8 Å and 41.2 Å, respectively) (**Figure S1**).

To investigate the association of the protein to the bilayer, we monitored the number of lipid atoms in contact with ASM during the simulation time (**Figure 2C**). We observed the spontaneous association of ASM in model 2 to the bilayer after 70 ns of simulation time. For model 1, we observed a similar overall trend for the three replicates, and the number of lipids in contact with ASM did not show marked fluctuations after 200 ns of simulation time. It has been suggested that the saposin domain of ASM plays a crucial role in the binding to the membrane ^11,12^. We thus monitored the residues within the saposin domain (residues W86-H169) that are in contact with the lipids in the simulations and compared our results to those obtained from a previous study ^12^. Our MD simulations presented partial insertion of the saposin domain in the bilayer, mostly involving helices H1, H2, and H4 (**Figure 2D-E**). Furthermore, when comparing the lipid contact profiles of model 2 with the three replicates of model 1, we observed that in model 2, there are fewer lipid atoms in contact with helix H2 and H3, indicating a less deep insertion of ASM into the lipid bilayer than observed in model 1 (**Figure 2D**). In light of these observations, we focused on the MD simulations from model 1 and its replicates for further analyses. The three replicates of model 1 all have consistent lipid contact profiles, showing a similar interaction with the bilayer. The partial insertion of the saposin domain of ASM into the lipid bilayer resembled a saposin-lipid binding mode previously described (i.e., named type I) (**Figure 2E**) ^12^.

The membrane interactions of ASM are particularly relevant when considering its proposed catalytic mechanism. The active site of the protein is not adjacent to the membrane, but rather distant from its surface (around 20 Å at the beginning of the simulation in model 1). Therefore, ASM, after binding the membrane, needs to recruit the lipids at its active site, where they undergo hydrolysis^11^. We thus investigated if lipids could form atomic contacts with the zinc ions, the catalytic (i.e., H321) and the zinc-coordinating residues (i.e., D208, H210, D280, N320, H427, H459, and H461), along with the suggested substrate-binding residues (i.e., N327, E390, and Y490) (**Figure 3 and S2**). We observed that starting from no lipids in their surrounding, all three replicates of model 1 consistently showed the substrate-binding residues N327, E390, and Y490 to form contacts with lipids, especially N327 and E390 (occurrence of lipid contact in the range 67.9-93.9% and 81.5-96.6% of MD simulation time) (**Figure 3A and S2**). N327, E390, and Y490 are located in the catalytic domain of ASM, in particular in the β3-α3, β5-α5 and β9-β10 loop, respectively, at the inner and hydrophobic concave surface, which faces the membrane (**Figure 3B**). In light of our results, N327, E390, and Y490 could play a role in the recruitment and orientation of the substrate lipids towards the catalytic site. We also observed lipid atoms in contact with the residues in the active site, especially involving H321, H459, and H461 in replicate 1 and 2 (occurrence in the range 31.4-91.5%, 54.6-82.1%, and 28.2-68.8% of MD simulation time, **Figure 3A and S2**). In terms of lipid species in contact with residues in the surroundings of the active sites, we observed different species, including sphingolipids (e.g., sphingomyelin) and glycerophospholipids (such as phosphatidylcholine, phosphatidylethanolamine, phosphatidylinositol, phosphatidylserine, and BMPs) (**Figure 3C and S2**). Our findings align with data from experimental micellar assay systems, which showed that recombinant ASM efficiently hydrolyzes sphingomyelins but also has hydrolytic capabilities, albeit with low efficiency, on a range of other membrane lipid species, including glycerophospholipids, lyso-glycerophospholipids, ceramide-1-phosphate, and BMPs ^9^. (**Figure 3C**). We noticed that BMPs, which are anionic lipids, form high-occurrence contacts with the zinc ions in replicates 2 and 3 (**Figure S2**), which might be due to an overstabilization of the ionic interaction between the metal ions and the anionic lipids in the MD force fields used for this study.

**Figure 3.**
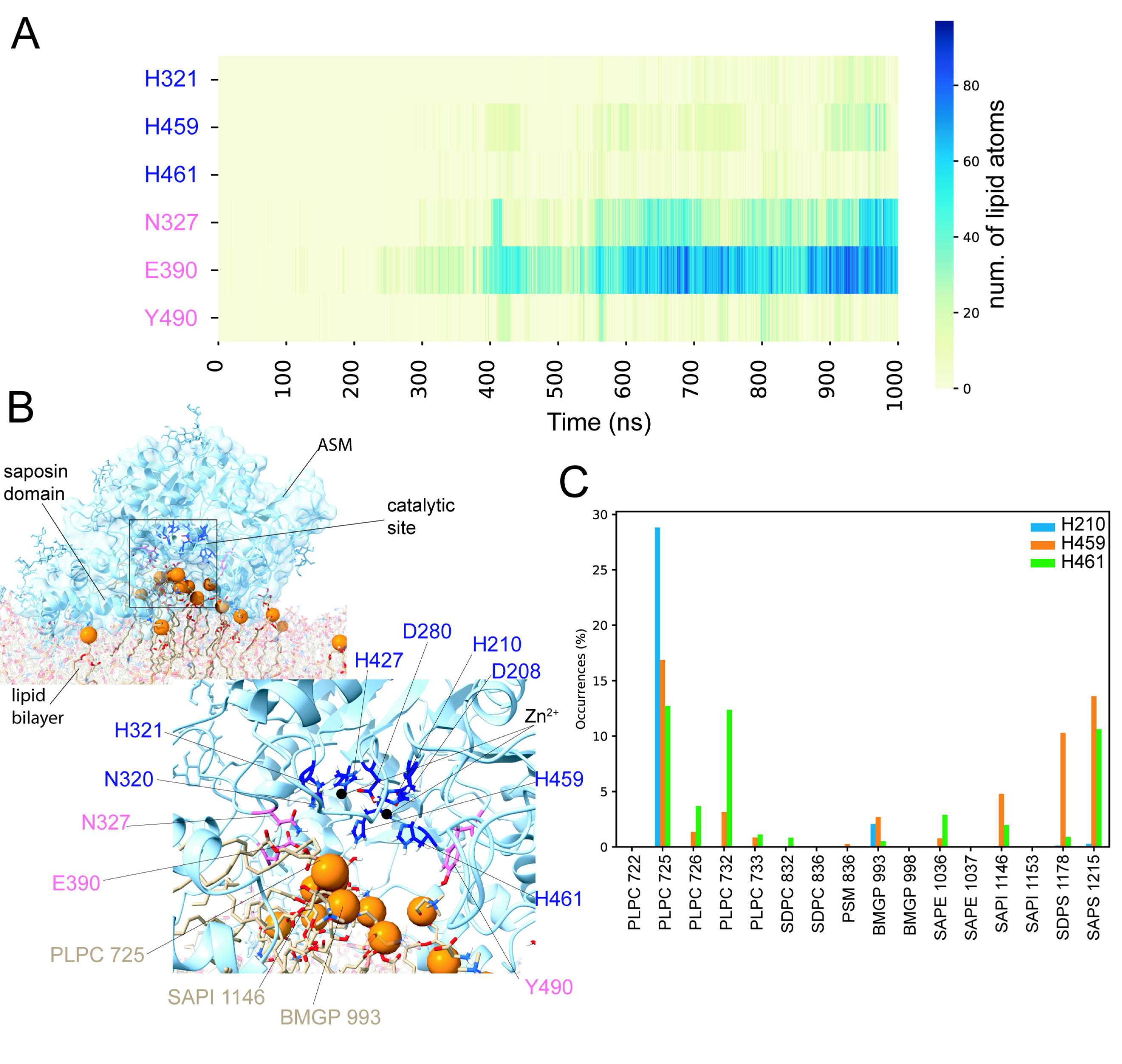
ASM recruits diverse lipid species at its catalytic site. (A) Heatmap showing the number of lipid atoms in contact with the substrate-binding (N327, E390, Y490), catalytic (H321), and zinc-coordinating residues (H459 and H461) of ASM during the replicate 1 of model 1. The other residues in the active site and the zinc ions featured contacts with the lipids for less than 20% of the MD replicate. (B) Visualization of the orientation of ASM relative to the lipid bilayer after 900 ns in replicate 1 of model 1. In the upper panel, ASM is shown as a blue cartoon, with lipids contacting the catalytic site residues depicted as light brown sticks, and their phosphate groups are highlighted as orange spheres. The lower panel highlights the orientation of lipids recruited at the active site (light brown) and the substrate-binding (pink), catalytic, and zinc-coordinating (dark blue) residues within the catalytic site of ASM. (C) Diversity of lipid species interacting with the active site residues of ASM during replicate 1 of model 1. The bar plot shows the occurrence of lipid species in forming contact with the residues of ASM. We observed various sphingophospholipids and glycerophospholipids, suggesting a broad lipid substrate specificity of ASM.

To further investigate the mechanism of lipid recruitment around the saposin domain and at the active site of ASM, we monitored the impact of protein adhesion on the membrane curvature (**Figure 4**). We observed that ASM induced a negative mean curvature in both bilayer leaflets in all the replicates of model 1, resulting in a dome-like shape beneath the active site (**Figure 4A-B**). The average local mean curvature under the protein active site reached values as low as -0.8 Å^−1^ for both the upper and lower leaflet, corresponding to a maximum average radius of curvature around 1.25 Å. Furthermore, for replicate 2 of model 1, ASM induced positive curvature in two regions of the bilayer, which correspond to the areas where the saposin domain inserts in the bilayer (**Figure 4B**). This observation suggests an effect on membrane curvature due to the insertion of the saposin domain into the bilayer in this replicate. Overall, the curvature induced by ASM to the bilayer in the region next to the saposin and beneath the catalytic site may facilitate the recruitment of lipids to the residues in the catalytic site.

**Figure 4.**
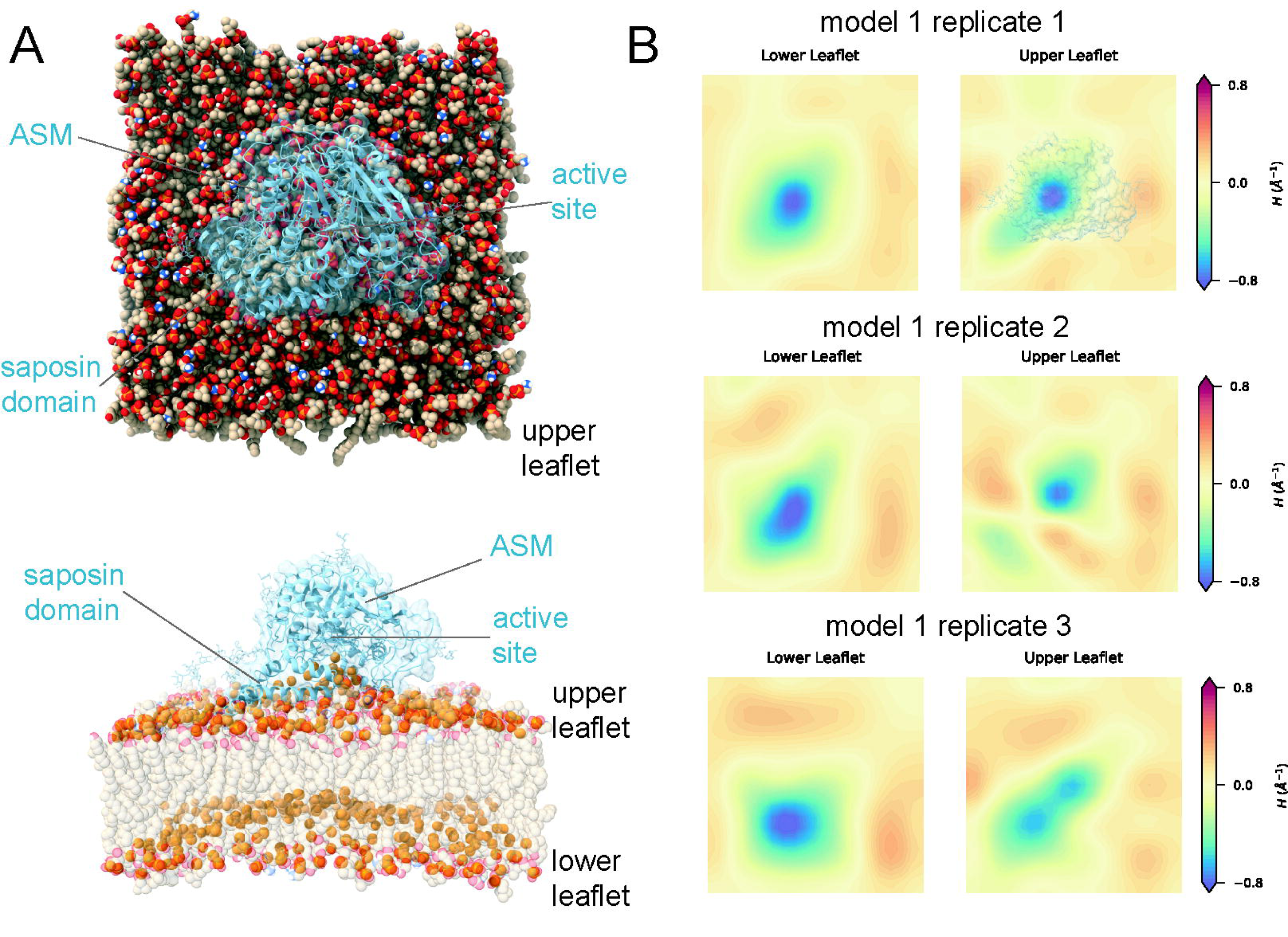
ASM induces membrane curvature in the bilayers. (A) Structure from replicate 1 of model 1, with an above view (upper panel) and side view (lower panel) of ASM. These views illustrate ASM inducing a negative mean curvature in both bilayer leaflets, resulting in a dome-like formation beneath its active site. (B) Average mean curvature of the lower (left panels) and upper (right panels) leaflets of the bilayer calculated for replicates 1-3 of model 1. Negative and positive mean curvature are indicated in blue and red, respectively. The average local mean curvature under the protein active site reached values as low as -0.8 Å^−1^. Replicate 2 shows positive curvature in two bilayer regions corresponding to insertion points of the saposin domain, suggesting a deeper insertion of the saposin domain in this replicate relative to the others.

### Parametrization of Cationic Amphiphilic Drugs (CADs): A Case Study with Ebastine

After investigating the binding of ASM to the lipid bilayer, we focused on studying the effects of CADs on the membrane biophysical properties, lipid-protein interactions, and the ASM structure.

To model CADs in MD simulations, it is necessary to compute the missing force-field parameters to describe the target molecules. Here, we used a workflow based on different computational approaches to define the force-field parameters for ebastine (**Figure 5A**) as a representative of cation amphiphilic drugs affecting ASM function at the lysosome (see Materials and Methods). In brief, we calculated the final set of force-field atomic charges and parameters for ebastine by analogy to existing parameters from an established force-field model, frequently used in MD simulations of membranes and small molecules ^39,40^ (**Figure 5A and Table 2**). Ebastine is a basic compound that contains a tertiary amine group, which is protonated at the lysosomal pH of ∼4.5-5.0. We thus generated parameters for ebastine for the protonation state that should be predominant at lysosomal pH (**Figure 5B**). We found that the force-field atomic charges and parameters we calculated for ebastine matched well with the existing force-field parameters, as shown by their low penalty scores (i.e., lower than 50) (**Table 2**). This consistency suggested that our parameters are reliable for use in our MD simulations of ebastine without needing further refinement. We used our parameters for ebastine to investigate at the molecular level its interactions and effects on membrane and ASM, such as curvature induction, by performing all-atom MD simulations in the presence of ebastine of i) ASM associated with the lysosomal-like bilayer (EBAH-ASM) and ii) the bilayer without the protein (EBAH) (**Figure 5C**).

**Figure 5.**
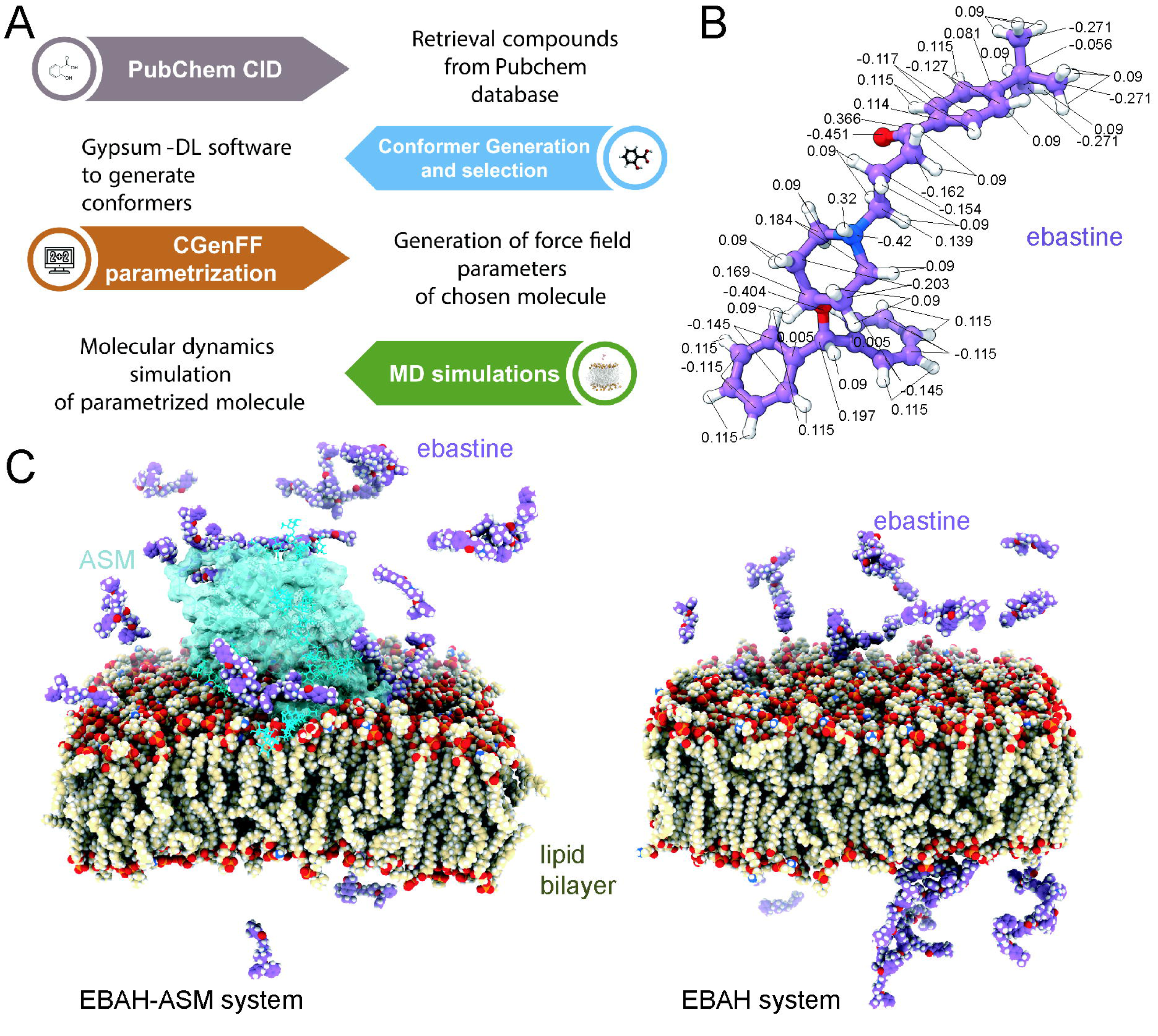
Force field parametrization of ebastine and design of MD simulations in membrane systems. (A) Outlines the computational workflow used to compute the force-field parameters for ebastine, employing computational approaches such as CHARMM General Force Field (CGenFF) for deriving parameters by analogy to existing parameters in the CHARMM36m/CGenFF4.6 force field. (B) Visualization of the protonation state of ebastine at lysosomal pH, showing the atomic charges obtained by CGenFF. (C) All-atom MD simulation designs of the two systems with ebastine: on the left, the combination of ebastine and ASM associated with the lipid bilayer (EBAH-ASM system); on the right, the lipid bilayer in the absence of the protein (EBAH system). Both systems included an ebastine/lipid ratio of 6%, a concentration previously shown to induce dose-dependent effects in bilayers with CAD-like molecules.

**Table 2.**
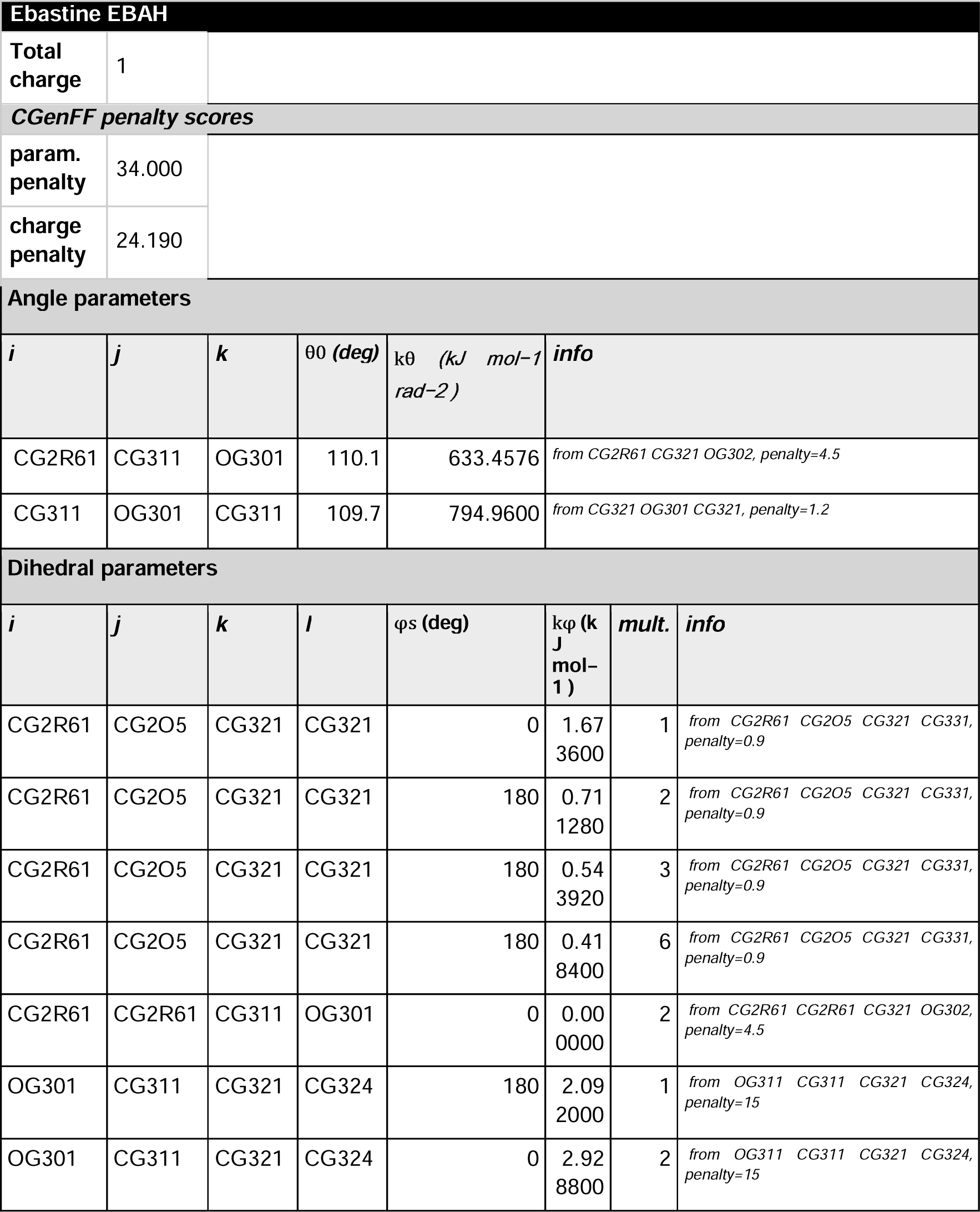
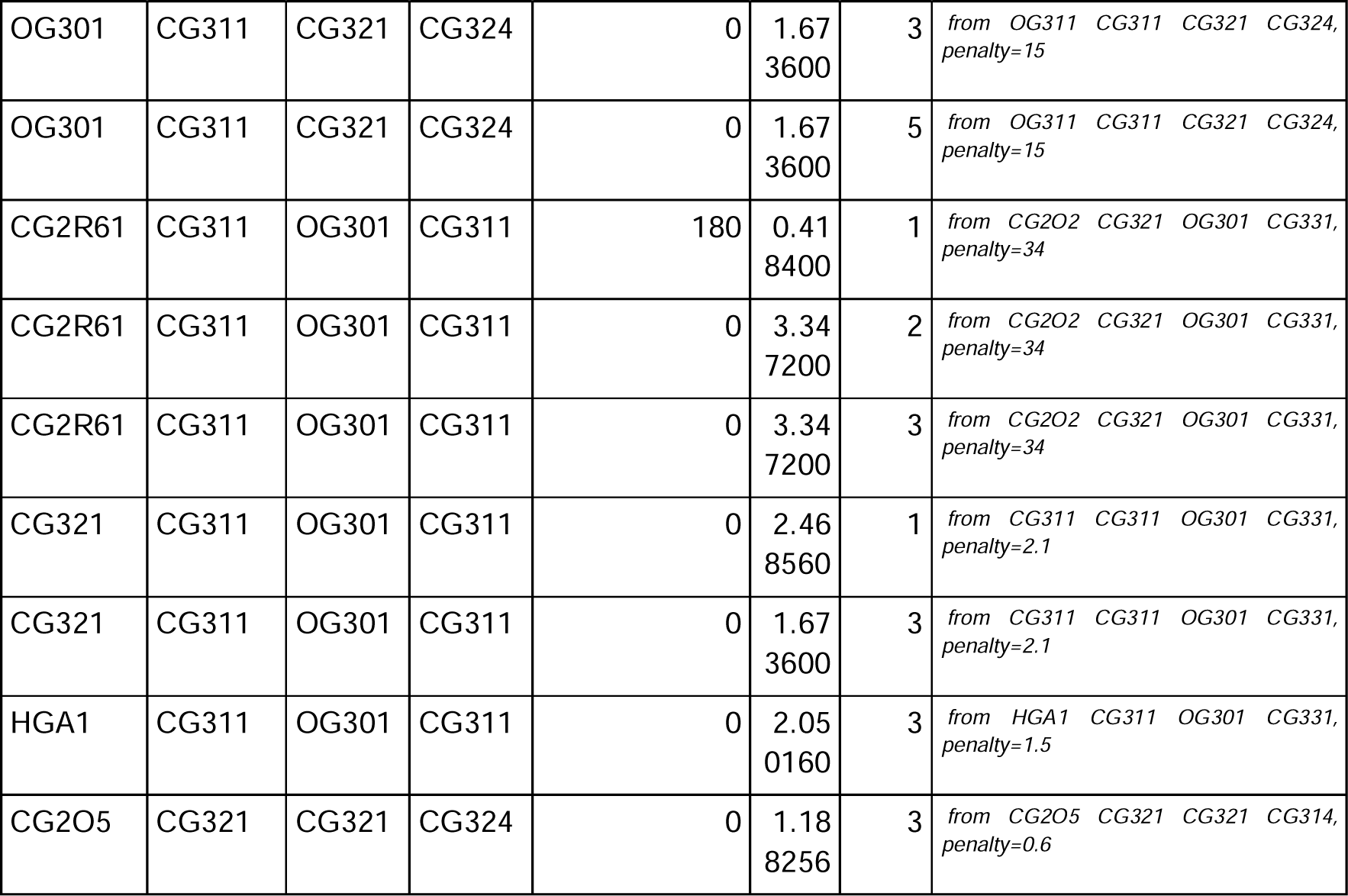
New force-field parameters for ebastine derived using CHARMM General Force Field (CGenFF) by analogy to existing parameters in the all-atom CHARMM36m CGenFF 4.6 force field.

We designed the MD simulations with ebastine following an approach recently applied to other CAD-like molecules ^38^. In this study, the authors performed simulations of lipid bilayers, including a membrane protein, with an overall system size comparable to our systems. They used two different CAD densities, i.e., a CAD/lipid ratio of 4.6% and 9.2% (equivalent to 24 and 48 CAD molecules, respectively). They observed that the amount of CAD molecules used in the simulation was enough to induce dose-dependent effects, such as alterations of membrane fluidity, curvature, thickness, and changes in the association of the protein with the membrane ^38^. We thus designed our systems with an ebastine/lipid ratio of 6% (equivalent to 37 ebastine molecules).

### How ebastine molecules alter the lysosomal membrane and interact with ASM

At first, we aimed to investigate the effect of ebastine on the lysosomal membrane itself. We thus estimated the number of ebastine atoms in contact with lipids in the MD simulations of the EBAH system (**Figure 6A**). We observed the spontaneous insertion of ebastine in the bilayer with more than 80% of ebastine atoms already in contact with the bilayer after 150 ns and nearly 100% after 300 ns (**Figure 6A**). On the other hand, in the EBAH-ASM system (**Figure 6B**), there was a slower and incomplete membrane insertion of ebastine. We observed that after 400 ns of simulation time, ∼70-80% of ebastine atoms are in contact with the bilayer (**Figure 6B**). Furthermore, the ebastine molecules integrated into the bilayer from either the upper or the lower leaflets in the EBAH and EBAH-ASM systems can be observed (**Supplementary videos S1-2**).

**Figure 6.**
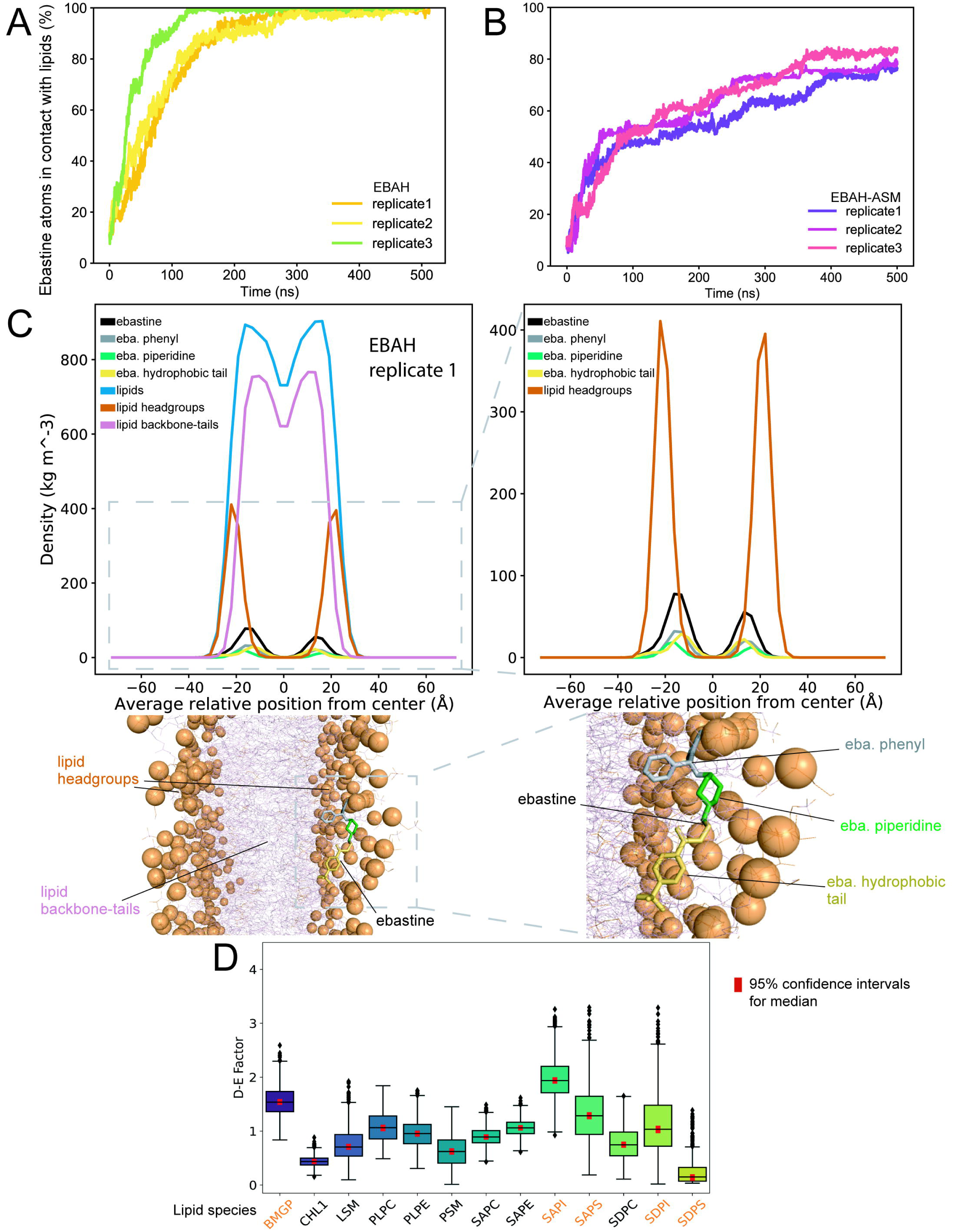
Ebastine interacts with a lysosome-like bilayer and inserts at the interface between lipid headgroups and tails. (A) Percentage of ebastine atoms in contact with the lipid bilayer over time in the EBAH replicates. We observed spontaneous insertion with over 80% of ebastine atoms in contact with lipids after 150 ns and full insertion after 300 ns. (B) Percentage of ebastine atoms in contact with the lipid bilayer over time in the EBAH-ASM replicates. We observed slower and not complete insertion of ebastine molecules in the bilayer in the presence of ASM with around 70-80% of ebastine atoms in contact with lipids after 400 ns (C) How ebastine and the main components of lipids are distributed within the thickness of the lipid bilayer. The left upper panel shows the mass density profiles of i) all ebastine atoms (black), ii) ebastine phenyl groups (grey), iii) ebastine piperidine groups (green), iv) ebastine hydrophobic tails (yellow), v) all lipid atoms (blue), vi) lipid headgroups (orange) and v) lipid backbone-tails (pink) along the axis perpendicular to the surface of the bilayer. For clarity, we show the results for replicate 1 of the EBAH system. The results for the other replicates are reported in Figure S3. The right upper panel highlights the mass density profiles of lipid headgroups and ebastine mass density profiles. The left lower panel shows a bilayer section with an ebastine molecule inserted in the membrane from replicate 1 of the EBAH system for illustrative purposes. Ebastine and lipids are shown as sticks, with the phosphate groups of the lipids highlighted as spheres. The right lower panel highlights the orientation of ebastine in the bilayer. The ebastine molecules insert at the interface between lipid headgroups and hydrophobic tails, with their hydrophobic tails and phenyl groups preferentially orienting towards the lipid tails and the piperidine ring towards the lipid headgroups. (D) The box plot displays the depletion-enrichment factor (D-E factor) for each lipid species in proximity to the piperidine group of ebastine, calculated for the replicate 1 of the EBAH system. Depletion-enrichment factor values above 1 suggest enrichment of the lipid species, whereas values below 1 indicate depletion. Anionic lipids are highlighted in orange. The red bars represent the 95% confidence intervals for the median of the depletion-enrichment factor values of each lipid species, calculated using bootstrapping analysis.

To disclose the mode of insertion and integration of ebastine into the membrane, we investigated how ebastine and the main components of lipids are distributed within the thickness of the lipid bilayer. We calculated the mass density profiles along the axis perpendicular to the surface of the lipid bilayer for each replicate of EBAH and EBAH-ASM (**Figure 6C and Figure S3**). We monitored the density of i) all ebastine atoms, ii) ebastine chemical groups, iii) all lipid atoms, iv) lipid headgroups, and v) lipid backbone-tail groups. This analysis was carried out on the final 200 ns of each simulation replicate. We chose this timeframe to ensure that nearly all the ebastine molecules had consistently integrated into the bilayer in the EBAH systems (**Figure 6A**). We observed that ebastine is generally inserted in the bilayer at the interface between lipid headgroups and their backbone-hydrophobic tails (**Figure 6C and Figure S3**). Furthermore, the analysis suggested that ebastine preferentially oriented with their hydrophobic tails and phenyl groups towards the lipid tails, while the piperidine ring, featuring the positively charged tertiary amine group, oriented more towards the lipid headgroups (**Figure 6C and Figure S3**). In the EBAH-ASM systems, we noted a comparable pattern for ebastine (**Figure S3**). However, in these systems, the incomplete integration of ebastine into the bilayer resulted in less defined density profiles.

We then estimated if the piperidine ring of ebastine, when inserted into the membrane, formed preferential interaction with any of the lipid species used in the modeling. In particular, we estimated the changes in the concentration of each lipid species around the piperidine ring, compared to their concentration in other regions of the membrane. This was achieved by calculating the depletion-enrichment factor ^41^ of each lipid species (**Figure 6D and Figure S4**). Similar to the density analysis mentioned earlier, this analysis was performed on the last 200 ns of each replicate of EBAH. This timeframe was chosen to ensure that nearly all ebastine molecules had integrated into the bilayer in the EBAH systems. Depletion-enrichment factor values above 1 indicate enrichment of the lipid species around ebastine, while values below 1 indicate depletion. Our analysis suggested a propensity for enrichment of anionic lipids in the vicinity of the piperidine ring of ebastine, including phosphatidylinositol, phosphatidylserine species, and BMPs (**Figure 6D and Figure S4**). In detail, we observed enrichment of BMPs (BMGP) and SAPI across all replicates of EBAH. These ebastine-lipid contacts could be driven by interactions between the positively-charged ebastine amine group and the lipid anionic headgroups, as previously proposed for another CAD-like molecule ^38,42^.

In the presence of ebastine and the absence of the protein, the lipid bilayer has an average lipid density of ∼ 14100 Å^3^ (**Figure S5**), thus slightly increased when compared to the MD simulations with ASM (∼ 13800 Å^-3^). We then calculated the average area per lipid and lipid bilayer thickness in the three replicates of the EBAH system (**Figure 7A and Figure S5**). We observed that the insertion of ebastine in the bilayer slightly increased its area per lipid (average 60 ± 0.13 Å^2^) and reduced the thickness (average 40 ± 0.4 Å) when compared to the reference MD simulations of a bilayer with the same lipid composition but in the absence of ebastine (average APL 56.6 Å^2^ and thickness 42.2 ± 0.2 Å) ^35^. The thickness profiles over time for EBAH, EBAH-ASM and ASM in absence of ebastine (model 1) simulations were similar (**Figure S5 and S6**), with only few conformations of EBAH-ASM at lower levels of thickness (< 39 Å). The area per lipid profile over time had a similar overall trend in the two systems but reached a plateau at lower values for EBAH-ASM (average area per lipid 5.66 ± 0.02 Å^2^) than EBAH **Figure S5**). The addition of ebastine did not affect the area per lipid of the bilayer when the protein is present (**Figure S6**). In absence of the protein, ebastine alone did not induce changes in bilayer curvature (**Figure 7B and Figure S7**). Moreover, ebastine did not have effects on the curvature induced by ASM itself on the bilayer in the EBAH-ASM systems compared to what observed with the ASM model 1 replicates (**Figure S8**).

**Figure 7.**
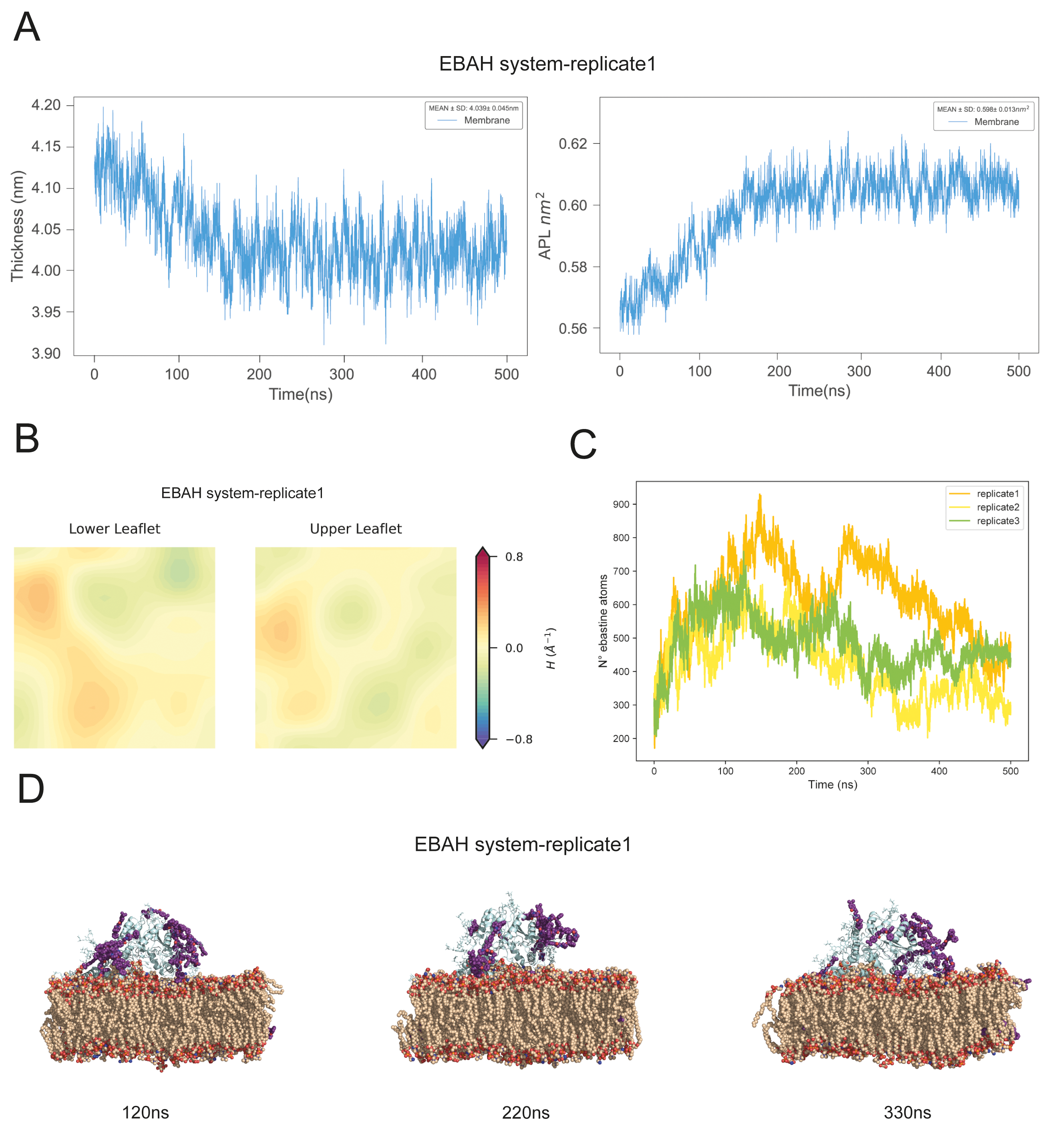
Membrane biophysical properties in presence of ebastine. (A) Average area per lipid (left panel) and lipid bilayer thickness (right panel) of replicate 1 of the EBAH system. Ebastine did not induce substantial changes in the thickness profiles of the EBAH, EBAH-ASM, and ASM model 1 simulations (**Figure S5-6**). Moreover, it induced only minor changes in both area per lipid and bilayer thickness compared to MD simulations in absence of ebastine^35^. (B) Average mean curvature of the lower (left panels) and upper (right panels) leaflets of the bilayer calculated for the replicate 1 of EBAH system. Negative and positive mean curvature are indicated in blue and red, respectively. Ebastine did not induce any specific strong positive or negative curvature in the bilayer. (C) Number of ebastine atoms in contact with the protein or the N-glycans over time in the EBAH-ASM replicates. (D) Three representative structures were extracted at 120 ns, 220 ns, and 330 ns from the first replicate of the EBAH-ASM simulation. ASM is depicted as a light blue cartoon, with lipids shown as spheres, and ebastine molecules as purple spheres. Ebastine formed high-occurrence contacts with ASM within the first 200 ns of simulation time, followed by a decrease as more ebastine molecules detached from the protein.

We evaluated the binding modes of ebastine into the ASM structure using a contact-based analysis combined with the analysis of solvent-exposed pockets of ASM (**Figures 7C-D and 8**) taken from our previous study ^37^. According to the previous analysis, pockets 1, 3, 6, 7, 15, and 22 matched with regions for ASM-membrane interactions, whereas pockets 5 and 12 included residues near the catalytic site. Ebastine established contact with the protein in all the MD replicates within the initial 100 ns (**Figure 7C-D, Supplementary Movie 3**).

**Figure 8.**
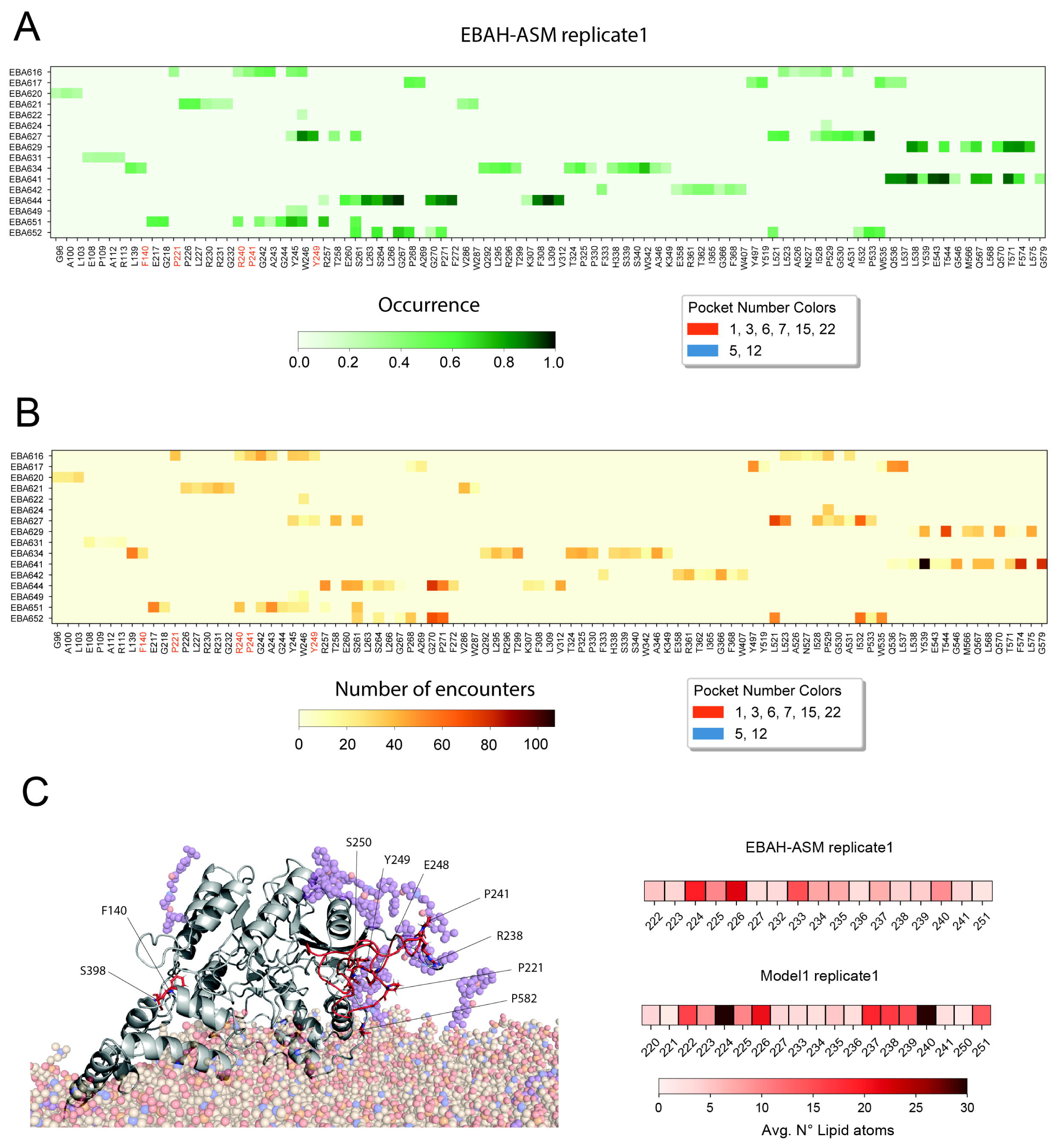
Ebastine interacts with ASM and interferes with its membrane binding. (A-B) The heatmaps show the occurrence (A) and the number of encounters (B) of the contacts between ebastine molecules and the residues of ASM in the replicate 1 of the EBAH-ASM system. The heatmaps report only ebastine-protein contacts occurring in at least 20% of the simulation frames. Residues located in the pockets which correspond to membrane-binding regions or those near the catalytic site are highlighted in dark orange and blue, respectively. (C) The left panel displays a representative structure extracted at 500 ns from replicate 1 of the EBAH-ASM simulation. The protein is depicted as a gray cartoon with the β1-α1 loop in red, while the lipids are shown as light brown spheres and ebastine molecules are highlighted as purple spheres. Residues within the pockets that make contacts with ebastine in any of the three replicates of EBAH-ASM are highlighted as red sticks. For clarity, *N*-glycans of the protein and all hydrogens are omitted. On the right panel, the heatmaps show the average number of lipid atoms in contact with the residues of the β1-α1 loop in the replicate 1 of EBAH-ASM (upper heatmap) and model1 (lower heatmap). The observed reduction in the average number of lipids in contact with the protein suggests that ebastine molecules, upon binding to ASM, interfere with the association to the membrane of the β1-α1 loop.

We estimated the occurrence (**Figure 8A-B**) and stability (**Supplementary Figure S9**) of the interaction of the different ebastine molecules with ASM in the three MD replicates, focusing on contacts formed at least in the 20% of the simulation frames. We observed that ebastine tends to bind to many different regions of the protein but with low occurrence and several events of formation/breaking of the interactions. In addition, we observed the tendency of forming interactions with some of the interfaces for membrane association and no interaction in the proximity of the active site. We observed that ebastine molecules interacted with residues in the β1-α1 loop (**Figure 8A-B**). This loop extends from the catalytic domain and localizes close to the H2, H3 and H2-H3 loop of the saposin domain^11^, potentially contributing to membrane binding. Interestingly, we identified that the ebastine molecules tend to reduce the lipid contacts of the residues in the β1-α1 loop (**Figure 8C**).

Our results suggest that ebastine in high concentration can interfere with the association of ASM with the lysosomal membrane, binding loops of the catalytic domain involved in its anchoring to the bilayer.

## Conclusions

In this study, we investigated the interaction of ASM with lysosomal membranes using microsecond all-atom molecular dynamics. We also modeled the effects of ebastine as a representative of cationic amphiphilic drugs on the lysosomal membrane and on ASM associated with the membrane. In our simulations, we accounted for the fully glycosylated form of ASM ^37^, and we designed a bilayer that mimics the lipid composition of typical lysosomal membranes from normal cells ^31,32^.

Overall, this work represented the first important step in developing a protocol to expand to other cationic amphiphilic drugs and membrane compositions. It also provided essential mechanistic insights into the structure of ASM associated with lysosomal membranes and the effects of ebastine. In detail, our results confirm the type I association between the saposin domain of ASM and the lysosomal membrane observed with coarse-grained models^12^. Our data also support the role of N327, E390, and Y490 in the recruitment and orientation of the substrate lipids to the active site. In addition, the simulations also show that the ASM association with the lysosomal membrane changes the membrane curvature and promotes a dome-like shape beneath the active site, which could facilitate the recruitment of lipids to the active site.

Furthermore, the presence of different lipid species, including sphingomyelins, glycerophospholipids, ceramide-1-phosphate, and BMP in the catalytic site of the enzyme during the simulation supports the promiscuous phospholipase activity of ASM over a wide range of lipid species.

Ebastine generally interferes with the bilayer at the interface between lipid headgroups and their backbone-hydrophobic tails. In particular, ebastine hydrophobic tails are oriented towards the lipid tails of the lysosomal membrane, and the ebastine piperidine ring interacts with the lipid headgroups. This is consistent with what was observed for other lysosomotropic drugs ^28^ and amphiphilic drugs or natural compounds ^43–46^. In addition, we observed an enrichment of anionic lipids near the piperidine ring of ebastine, including phosphatidylinositol, phosphatidylserine species, and BMPs. Notably, it has been proposed that another CAD-like molecule, upon intercalation into membrane models, formed contact with phosphatidylserine species, mainly driven by electrostatic interactions with the lipid headgroups, and restricted their lateral diffusion, making them less available for the binding with peripheral membrane proteins ^38,42^.

According to the timescale and the membrane size and composition used in this study, we only observed minor changes in area per lipid and thickness induced by ebastine and no changes in other membrane biophysical properties, which might require additional sampling or larger membrane constructs. Other factors could be related to the ebastine/ratio used in this study. A future step should include additional simulations changing the ratio to observe dose-dependent effects, similar to what was done in other works ^23,38^.

Cation amphiphilic drugs are expected to cause alterations primarily to membranes of lysosomes from cancer cells, which often have alterations in lysosomal compositions with respect to lysosomes from normal cells, such as in their sphingolipid metabolism ^6,30^. Thus, on one side, our results could also be interpreted as a genuine pattern of mild changes induced by ebastine on normal lysosomal membranes. This is prompted by using a bilayer composition resembling one of the lysosomes in a non-transformed form. A future natural step would be to include data from lysosomal lipid profiling in cancer cells using lipidomics and exploit techniques for organelle-level lipidomics ^31,32^ from cancer cells before and after CAD treatment, such as lipidomics data from leukemia cell lines ^30^.

At the best of our knowledge, it is still unclear if CADs can directly bind to ASM and other lysosomal hydrolases ^14^. Our results show that ebastine can directly interfere with ASM, altering loops of the catalytic domain that serve as anchors for the interaction with the membrane. This can be the first step to destabilize ASM association with the lysosomal membrane. We did not expect to observe a detachment of ASM from the lipid bilayer in the microsecond timescale sampled by the unbiased MD simulations. Larger sampling of the conformational space and enhanced sampling approaches, along with higher CAD concentrations, are needed to probe if molecules as ebastine could induce ASM dissociation from the lysosomal membrane.

In addition, the ASM-EBAH simulations presented in this study support a model of action in which CADs could compete with the lysosomal hydrolases, such as ASM, in the binding to anionic lipids and neutralize the negative charges on the surface of the intraluminal lysosomal vesicles ^6,14^.

More broadly, our work provides a computational approach that can be applied to other CADs using the parameterization protocol provided here and the strategies for designing the initial configuration for MD, together with the tools from the LipidDyn package ^47^.

## Methods

### Design of lipid bilayers

We used the CHARMM-GUI Membrane Builder tool ^48^ to design a lipid bilayer with a composition resembling a recent model of the membrane of mammalian lysosomes ^35^ (**Table 1**). We build a symmetric lipid bilayer of 125 and 125 Å in the x and y dimensions, respectively. The lipid bilayer included a total of 306 lipids per leaflet. We designed two systems with different starting orientations of ASM to the lipid bilayer. In the first system, we modeled ASM as already localized on top of the lipid bilayer by using the initial configuration previously published ^37^. In the second system, we avoided imposing extensive starting contacts between ASM and the lipids using an approach similar to the one used for other peripheral membrane proteins ^38^. Starting from the orientation of ASM in the first system, we used the CHARMM-GUI Membrane Builder tool to translate the protein of 15 Å along the z-axis and rotate it by 15 and -10 degrees on the x and y-axis, respectively. We solvated the systems in a rectangular box of water molecules of 157 Å and 167 Å in the z dimension for the first and second systems, including around 320 and 340 water molecules per lipid, respectively. We used the lipid2MD tool available in LipidDyn (https://github.com/ELELAB/LipidDyn)^47^ to analyze the lipidomics datasets of lysosomes and match each measured lipid species with the corresponding parameters available in CHARMM36m force field ^39^. We used these two systems, hereinafter referred to as model 1 and model 2 for the former and latter modeling strategy, respectively, as starting structures to perform all-atom molecular dynamics simulations.

Furthermore, we used the last structure of the trajectory of model 1 to design the starting structure for MD simulations, including ASM and ebastine. We started from model 1 in light of the analysis of contacts between the protein residues and the lipids, which pointed out a deeper insertion of the saposin domain into the membrane than what was observed for the simulations starting from model 2. We verified that the last structure of replicate 1 of MD simulations for model 1 represented the entire trajectory through structural clustering with the GROMOS algorithm ^49^ on the Cα-atoms RMSD matrix.

### MD simulations

We used the MD simulation of model 1 previously published ^37^ and performed two additional all-atom MD simulations of one-μs each for ASM embedded in a lipid bilayer with a mammalian “lysosomal-like” composition ^35^ with the CHARMM36m force field ^39^. Moreover, we carried out one additional simulation using model 2 (**Table 1**).

To study the effects induced by ebastine, we also performed additional replicates of 500 ns for the same bilayer, including ebastine (EBAH), as well as ebastine and ASM (EBAH-ASM) with the CHARMM36m force field ^39^. To this goal, we used the force field parameters for ebastine obtained as described in the section above (**Table 2**). For the system with ebastine and ASM attached to the membrane, we used the last frame of replicate 1 of model 1 as a starting structure. Following a previously published approach ^38^, we included ebastine in the two systems at a density of 6% ebastine/lipid ratio as discussed in the Result Section. We used the *insert-molecules* tool of GROMACS to randomly insert the ebastine molecules into the box of water molecules.

We minimized and equilibrated the systems following the standard CHARMM-GUI Membrane Builder protocol. The protocol consists of multiple equilibration steps, which gradually release positional and dihedral restraints, described by harmonic restraints, applied to the protein and membrane. In addition, we performed a final equilibration, releasing the restraints mentioned above. Firstly, we performed 10000 steps of minimization by the steepest descent method. Following, two short simulations in the canonical ensemble (i.e., NVT) of 250 ps with an integration step of one fs, using Berendsen thermostat ^50^ with a coupling constant of one ps, to reach the desired temperature of 310K. The two thermalization phases were followed by four pressurization steps constituted by 250 ps, 500 ps, 500 ps, and five ns with an integration step of one, two, two, and two fs, respectively. Regarding all the steps, the pressure was controlled semi-isotropically using Berendsen barostat ^50^, with a time constant of five ps. The final equilibration to finalize the systems lasted five ns with an integration step of two fs.

The productive simulations were performed using a time step of two fs in an NPT ensemble, employing the Nose-Hoover thermostat ^51^ Parrinello-Rahman barostat ^52,53^ with time constant of one ps and five ps, respectively. In addition, the LINCS algorithm ^54^ was used to constrain heavy-atom bonds and the cutoff for both Van Der Waals and Coulomb interactions was set to 1.2 nm, as well as the particle-mesh Ewald scheme with a 0.12 nm grid spacing ^55,56^.

### Parametrization of CHARMM36 force field for ebastine using CGenFF

We retrieved the two-dimensional (2D) chemical representation of ebastine as SDF format files from the PubChem database (PubChem entry https://pubchem.ncbi.nlm.nih.gov/compound/3191, CID:3191) ^57^. We then used GYPSUM-DL v.1.1.5 ^58^ to convert the 2D representation of ebastine into 3D models and define the appropriate protonation states at the selected pH. For the calculation with GYPSUM-DL, we used 4.5-5.0 and 7.0-7.5 as pH ranges to be considered, and we used default values for the other parameters. For each pH range, we generated ten conformers for ebastine. We then selected two conformers for the next parametrization steps: i) one with the protonation state that should be predominant at pH 4.5-5.0 (i.e., lysosomal pH) ^59^ and ii) its unionized form presents at pH 7.0-7.5 (i.e., cytosol pH) ^60^. We parametrized both the protonation states of ebastine to check that the generated parameters and topologies were consistent with each other. We used open babel 3.1.1 ^61^ to convert the output files from GYPSUM-DL into mol2 format files. We employed the stand-alone version of the CHARMM General Force Field (CGenFF) program v. 2.5.1 (SilcsBio)^62^. CGenFF automatizes the generation of force field parameters by defining molecule topology, atom typing, atomic charge assignment, and parameters based on analogy to existing parametrizations in the target force field. CGenFF provides a penalty score for each atomic charge and parameter assigned, evaluating their analogy to existing parameters (i.e., low penalty scores mean good analogy). We calculated the parameters for ebastine using as targets the already available parameters in the all-atom CGenFF 4.6 ^40^. We verified that in the set of parameters we developed for ebastine, no terms with penalty scores greater than 50 would require further optimization, such as by ab initio calculations. We then converted the novel parameters from the format compatible with CHARMM software to the one of GROMACS (see Python script in GitHub repository). Finally, we included the topologies and the parameters for ebastine in the version for GROMACS 2022 of CHARMM36m ^39^ and CGenFF 4.6 ^40^. All the produced topologies and parameters for ebastine and a version of the CHARMM36m/CGenFF force field are available in the OSF repository associated with the publication.

### Analysis of MD simulations

We used LipidDyn ^47^ to calculate i) thickness, ii) area per lipid (APL), and iii) a two-dimensional (2D) lipid density map. In addition, we employed a new module of LipidDyn developed to calculate the mean (H) curvature of the bilayer. The implementation of the bilayer curvature was officially included in the LipidDyn GitHub repository on 17/01/2023 (https://github.com/ELELAB/LipidDyn/pull/121). LipidDyn uses the *LeafletFinder* class of MDAnalysis ^63^ to identify the leaflet of the bilayer and the lipids belonging to them, considering their headgroup for representative atoms of each lipid molecule. LipidDyn estimates the bilayer thickness as the distance vector between neighborhood-averaged coordinates of each lipid and its neighbors in the opposite leaflet, using a cut-off distance of 60 Å. Furthermore, APL is calculated by a neighbor search of each lipid and computation of a Voronoi tessellation. LipidDyn provides not only lipid density maps, using a density calculation algorithm analogous to the *densmap* tool of GROMACS, but also an estimate of the average lipid density during the simulation time^64^. LipidDyn also employs a reimplemented version of the *MembraneCurvature* tool, available in MDAnalysis [https://github.com/MDAnalysis/membrane-curvature/], to derive the membrane surface using lipid headgroups as reference atoms and then estimate mean curvature. For the comparison between EBAH-ASM and model1, the membrane biophysical properties were computed over the last 200 ns of the replicates.

We used a protocol previously applied to other cases to investigate the association of ASM with the bilayer and estimate the contacts between protein residues and lipids ^65^. In particular, we calculated the number of lipid atoms contained within a spherical surround with a radius of 6 Å around every protein atom during i) all the trajectory time and ii) over the last 500 ns of the trajectory. We also analyzed, in a similar manner, the lipids in contact with the zinc ions or with a selection of residues in the proximity of the catalytic site. In the analysis, we did not consider residues for which each atom featured contacts with the lipids for less than 20% of the MD frames.

To investigate the interaction mode of the ebastine molecules with the lipid bilayer, we employed the *density* tool from GROMACS to calculate mass density profiles along the z-axis of the simulation box. These profiles were generated for all atoms of ebastine and the lipids and specifically for the heavy atoms of the piperidine group, hydrophobic tail, and phenyl group of ebastine. Additionally, we calculated the profiles for the heavy atoms of the lipid headgroups and their backbone tails. Furthermore, we calculated the contacts between the ebastine and the membrane, i.e., we calculated the number of lipid atoms of each species in the vicinity of each ebastine molecule. Specifically, we used the GROMACS tools to calculate the number of lipid atoms of each lipid species in the vicinity of all the heavy atoms of the piperidine group of ebastine, using a distance cutoff of 6 Å. We then calculated the depletion-enrichment factor ^41^ around the piperidine group of ebastine of each lipid species as

depletion-enrichment factor of lipid species *X*:

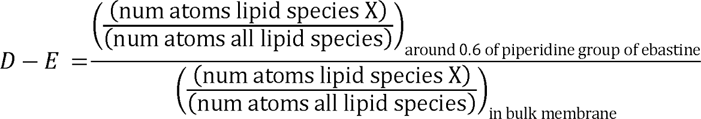

Depletion-enrichment factor values above 1 indicate enrichment of the lipid species, while values below 1 indicate depletion. We conducted bootstrap analysis to determine the 95% confidence intervals for the median of the depletion-enrichment factor values of each lipid species, setting the number of resamples to 1000 (see GitHub for code to reproduce the analysis).

We calculated pairwise atomic contacts among the heavy atoms of ebastine and the ones of the protein using CONtact ANalysis (CONAN) ^66^ to analyze interactions between ASM and ebastine molecules in the MD replicates. We applied a *r_cut_* cutoff of 10 Å, *r_inter,_* and *r_high-inter_* values of 4.5 Å. Residues with less than 20% contact with ebastine in the simulation frames were excluded from further analysis due to their low occurrence. We then combined this data with information on pocket residues of ASM as reported in ^37^ to evaluate if ebastine could contact sites for ASM association with the lysosomal membrane.

## Author Contributions (CRediT Classification)

*Conceputalization*: EP *Data Curation*: KM, SS, ML *Formal Analysis*: SS, ML *Funding Acquistion:* MJ, EP *Investigation*: SS, ML, EP *Methodology*: SS, ML *Project administration:* EP *Resources*: MJ, EP *Supervision*: ML, EP *Validation*: SS, ML *Visualization*: SS, ML *Writing – Original Draft:* SS, ML, EP. *Writing – Review and Editing*: All the co-authors.

## Acknowledgments

Our research has been supported by Danmarks Grundforskningsfond (DNRF125) and Novo Nordisk Fonden Bioscience and Basic Biomedicine (NNF20OC0065262) to the E.P. group. Part of the MD simulations have been performed thanks to the Danish HPC Infrastructure Computerome2 access. We would like to thank Mikkel Rohde for the fruitful discussions and comments.

